# *Atf6*^-/-^ mouse photoreceptors exhibit novel ciliary rootlet defect

**DOI:** 10.64898/2026.02.18.701862

**Authors:** Allyssa Bradley, Kristen Haggerty, Eun-Jin Lee, Michael A. Robichaux, Jonathan H. Lin

## Abstract

ATF6 is a regulator of the Unfolded Protein Response that maintains cellular homeostasis during ER stress. In patients, *ATF6* mutations cause photoreceptor dystrophy and sensorineural hearing loss. *Atf6*^*-/-*^ mice develop progressive hearing loss with stereocilia disorganization and mild retinal dysfunction, suggesting that ATF6 loss may impair the structural integrity of sensory cells. To test this possibility, we analyzed the retinal ultrastructure of *Atf6*^*-/-*^ mouse photoreceptors using transmission electron microscopy and identified a novel defect in which the ciliary rootlet is unbundled, disorganized, and possibly detached from the basal body. These findings demonstrate that ATF6 is essential for maintaining the structural organization of the photoreceptor ciliary apparatus, linking ER proteostasis to cytoskeletal integrity and providing a potential mechanistic basis for the progressive degeneration of photoreceptor outer segments and stereocilia observed in ATF6-deficient patients.

## Introduction

Activating Transcription Factor 6 (ATF6) is a protein involved in the Unfolded Protein Response that regulates ER stress and maintains cellular homeostasis (Haze et al., 1999; Walter and Ron, 2011; Hillary and Fitzgerald, 2018). In humans, biallelic *ATF6* mutations cause inherited retinal degenerations (IRD) including cone-rod dystrophy and achromatopsia with severe photoreceptor outer segment loss and foveal hypoplasia (Kohl et al., 2015; Kroeger et al., 2021; Skorczyk-Werner et al., 2017), and some of these IRD individuals also develop progressive sensorineural hearing loss (Lee et al., 2025). In mice, *Atf6* deletion leads to marked hearing deficits with stereocilia disorganization and mild late-stage visual decline (Kohl et al., 2015; Lee et al., 2025). Moreover, *Atf6*^*-/-*^ retinas show accelerated degeneration in the P23H rhodopsin model yet reduced pathological neovascularization in oxygen-induced retinopathy (Lee et al., 2021; Bradley et al., 2025). These divergent outcomes suggest cell-type-specific roles for ATF6 in proteostasis and stress adaptation.

Photoreceptor integrity relies on ER-to-cilium trafficking of outer segment proteins and on structurally supportive cytoskeletal elements (Young, 1967; Wensel et al., 2021). Because ATF6 regulates numerous ER chaperones and structural factors involved in secretory trafficking (Haze et al., 1999; Wu et al., 2007; Yoshida et al., 2000), its absence could gradually impair the folding, trafficking, or stability of proteins required for proper ciliary organization. We hypothesized that ATF6 deficiency leads to ultrastructural defects in the photoreceptor ciliary apparatus. To test this, we performed transmission electron microscopy (TEM) on six-month-old *Atf6*^*-/-*^ retinas. We discovered disorganization of the ciliary rootlet, suggesting that impaired ER proteostasis compromises photoreceptor cytoskeletal integrity which may underlie the gradual retinal vulnerability in *Atf6*^*-/-*^ mice.

## Materials and Methods

### Animals

Wildtype (*Atf6*^*+/+*^) and *Atf6*^*-/-*^ C57BL 6/J mice of both sexes at 6 months of age were used. All animal procedures were approved by the Institutional Animal Care and Use Committee of West Virginia University and conducted in accordance with the Association for Research in Vision and Ophthalmology (ARVO) Statement for the Use of Animals in Ophthalmic and Vision Research. *Atf6*^*-/-*^ mice were generated by the Kaufman laboratory (Wu et al., 2007) and provided previously to the Dr. Jonathan Lin laboratory by Dr. Randal J. Kaufman (Lee et al., 2021; Bradley et al., 2025; Lee et al., 2025).

### TEM

Retinal ultrastructure was analyzed as described previously (Thompson et al., 2025). Briefly, mouse eyes were enucleated, the anterior segments removed, and eyecups immersion-fixed in ice-cold 4% paraformaldehyde and 2.5% glutaraldehyde in 1×PBS (pH 7.4) for 12–16 h at 4 °C. Fixed retinas were dissected, rinsed in 100 mM cacodylate buffer (pH 7.4), and post-fixed in 1% osmium tetroxide plus 1.5% potassium ferrocyanide for 1 h at 4°C. Samples were stained in 2% uranyl acetate, dehydrated through graded ethanol and acetone, and embedded in Eponate 12 resin (Ted Pella). Ultrathin sections (70 nm) were cut on a Leica UCT ultramicrotome, stained with 1.2% uranyl acetate and 3% lead citrate. Images were obtained using a JEOL JEM-1010 or JEM-1400 transmission electron microscope.

## Results

### Ultrastructure reveals defective rootlet in Atf6^-/-^ photoreceptor inner segments

The photoreceptor ciliary rootlet is a striated cytoskeletal structure composed primarily of rootletin, which polymerizes into parallel filament bundles extending from the basal body deep into the inner segment (Yang et al., 2002). Because *Atf6*^*-/-*^ mice exhibit dysregulation of actin cytoskeletal genes and stereocilia disorganization in cochlear hair cells (Lee et al., 2025), we examined whether *Atf6* loss similarly affects photoreceptor cytoskeletal organization. TEM of six-month-old *Atf6*^*+/+*^ retinas revealed compact, well-aligned rootlet fibers forming a single tight bundle descending from the basal body into the inner segment (Fig. 1, arrowhead). In contrast, *Atf6*^*-/-*^ photoreceptors displayed marked rootlet disorganization: the fibers appeared unbundled, with multiple striated filaments present within individual inner segments (Fig. 1, arrow). Some fibers were misoriented or extended aberrantly past the basal body toward the apical direction, suggesting loss of proper anchoring (Fig. 1, bottom, white arrow). These ultrastructural abnormalities indicate that *Atf6* is required for maintaining the organization and anchoring of the ciliary rootlet in photoreceptors.

**Fig. 1.**
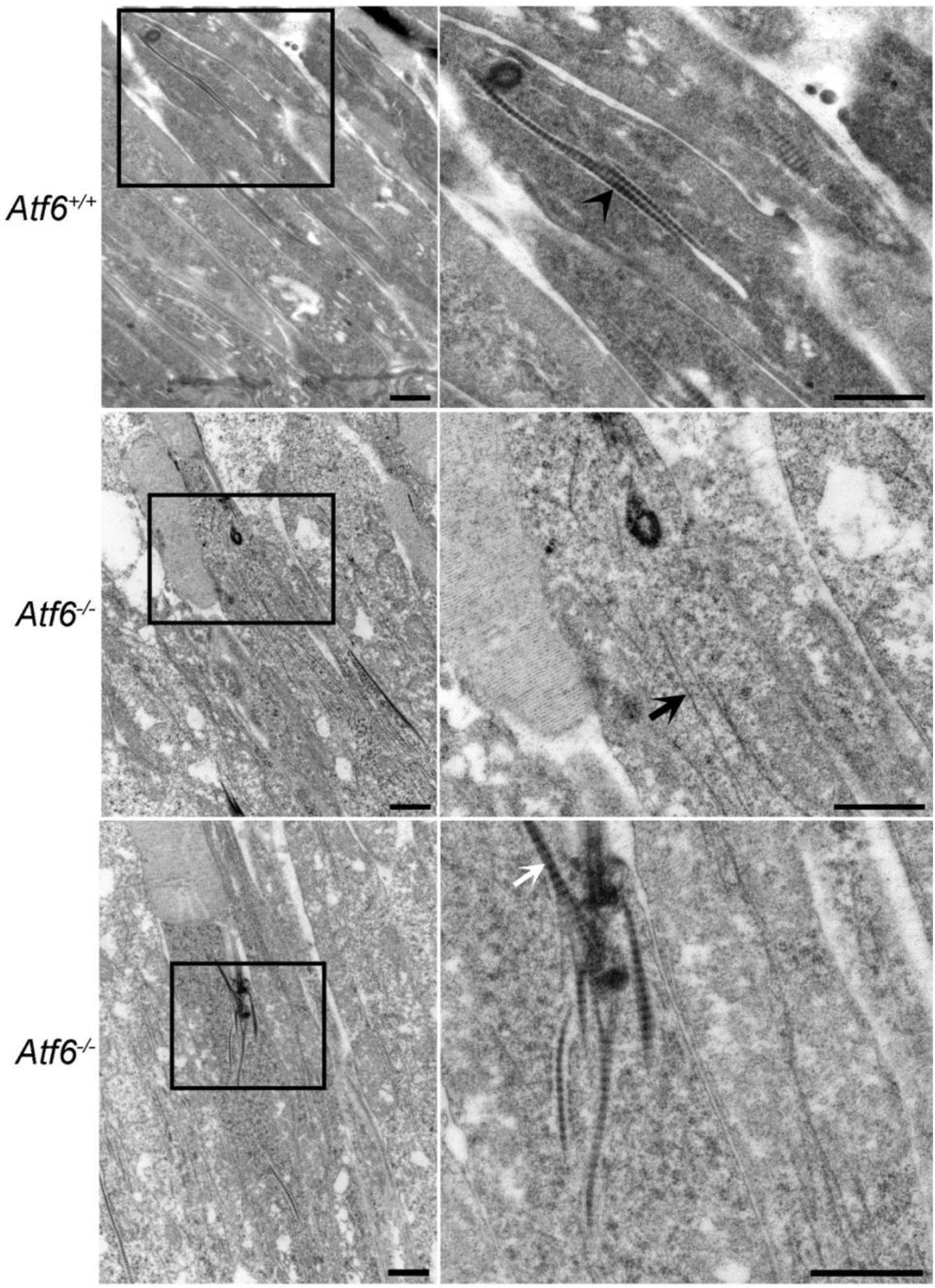
Disorganization of the photoreceptor rootlet in *Atf6*^−*/*−^ retinas. Transmission electron microscopy (TEM) images of photoreceptor inner segments from *Atf6*^*-/-*^ (top) and *Atf6*^*-/-*^ (middle and bottom) retinas. In wild-type photoreceptors, the rootlets appear tightly bundled (black arrowhead). In contrast, *Atf6*^*-/-*^ photoreceptors display unbundled rootlets (black arrow) and occasionally show misoriented rootlets extending upward from the basal body (white arrow). Insets show higher magnifications of boxed regions. Scale bars, 1 μm.

### Basal body disorganization accompanies rootlet detachment in Atf6^-/-^ photoreceptors

Given the loss of rootlet anchoring in *Atf6*^*-/-*^ photoreceptors, we next examined whether the basal body itself exhibited structural alterations. In wild-type cells, the basal body appeared compact and electron-dense, forming a continuous interface with the rootlet (Fig. 2, left, black arrowhead). In *Atf6*^*-/-*^ photoreceptors, the basal body region appeared less compact, and the rootlet appeared partially detached (black arrow). The anchoring zone contained irregular, electron-dense aggregates (dark circles, white arrowhead) interspersed with large, electron-lucent vesicle-like profiles (white circles, white arrow), features not present in controls. These findings indicate that Atf6 deficiency leads to disorganization of the basal body rootlet complex.

**Fig. 2.**
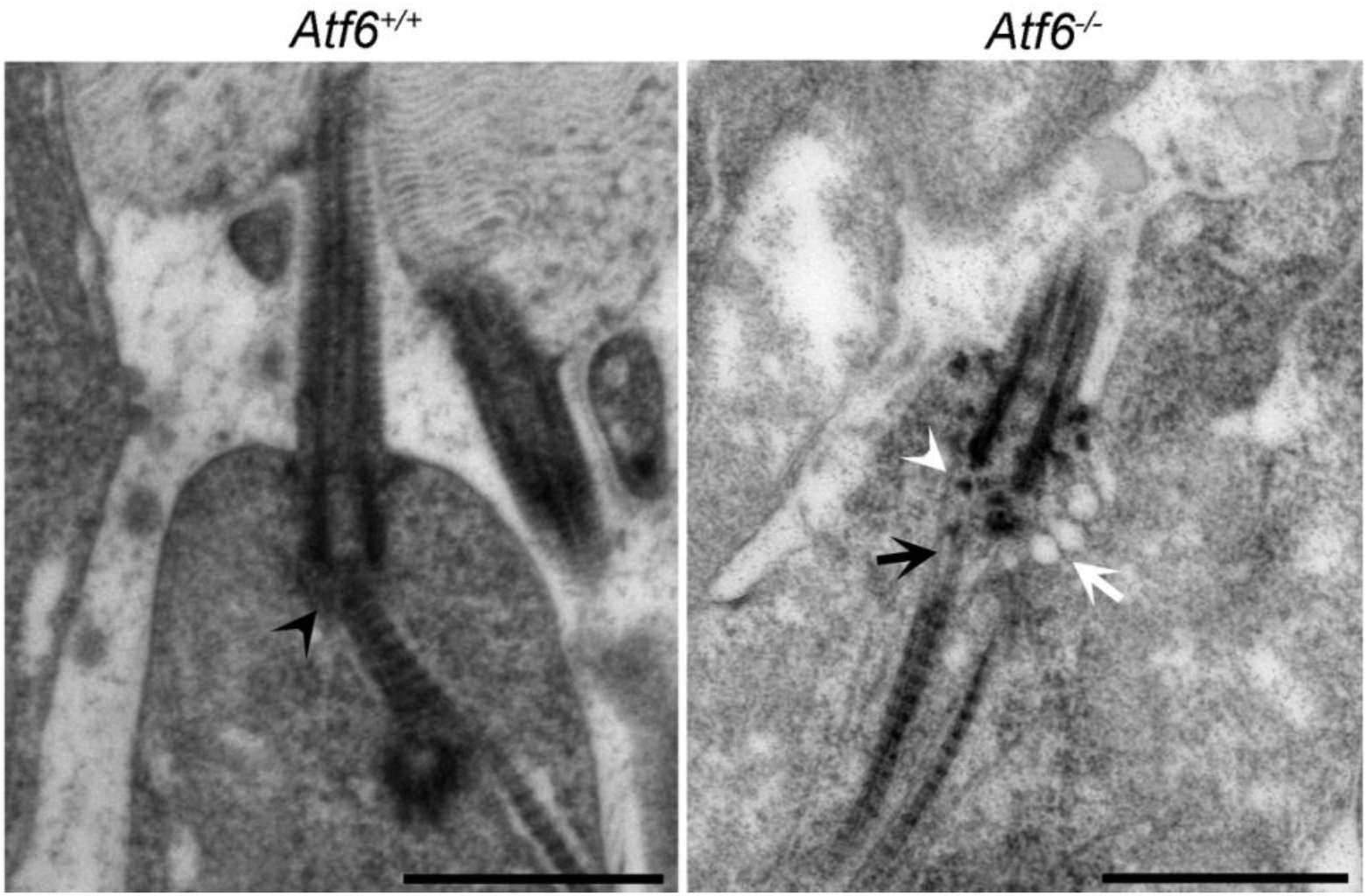
Basal body disorganization in Atf6^*-/-*^ photoreceptor inner segments. Transmission electron microscopy (TEM) images showing the basal body region of photoreceptor inner segments in *Atf6*^+/+^ (left) and *Atf6*^*-/-*^ (right) retinas. In wild type, the basal body is compact and connected to tightly bundled rootlet fibers (black arrowhead). In *Atf6*^−/−^ photoreceptors, the anchoring zone shows electron-dense aggregates (white arrowhead) and large, electron-lucent vesicle-like structures (white arrow). Scale bars, 1 μm.

## Discussion

Here we identify a novel rootlet defect in the *Atf6*^-/-^ mouse photoreceptor. Disruption of ciliary structures is a common cause of combined vision and hearing loss syndromes (Adams et al., 2007). We previously reported that *Atf6*^*-/-*^ mouse cochlear hair cell stereocilia are also structurally defective, and patients with ATF6 mutations are known to experience vision and hearing loss (Kohl et al., 2015; Lee et al., 2025). Together, these findings suggest that ATF6 deficiency may contribute to a form of sensory ciliopathy.

ER stress response pathways are known to play a role in vision and hearing loss (Li et al., 2024; Chan et al., 2024). Sensory cell types with cilia such as cochlear hair cells and photoreceptors are particularly sensitive to protein damage due to light and noise stimulus and oxidative stress from their high metabolic needs (Yang et al., 2008; Jongkamonwiwat et al., 2020). Our findings of structural defects in both the photoreceptor rootlet and cochlear stereocilia indicate that the ATF6 is critical for maintaining cytoskeletal stability in these highly specialized cells. Actin, a major component of both the stereocilium and rootlet, is known to be sensitive to oxidative stress, and the ER-mitochondria interface supports actin polymerization and actin-regulating signaling (Ong et al., 2020).

The rootlet is important for photoreceptor structural integrity. *Crocc*^*-/-*^ photoreceptors are extremely fragile in mouse retinas, which likely contributes to the mild retinal degeneration by 18 months, paralleling the *Atf6*^*-/-*^ mouse phenotype (Yang et al., 2005). In contrast, patients with ATF6 mutations exhibit severe cone-rod dystrophy or achromatopsia, likely reflecting interspecies differences in photoreceptor architecture (Wensel et al., 2021). The greater structural similarity between human and mouse cochlear stereocilia may also explain why *Atf6*^*-/-*^ mice and Usher syndrome mouse models display prominent hearing loss but minor or no retinal degeneration (Ohlemiller, 2019).

The differences between mouse and human photoreceptors make the *Atf6*^*-/-*^ model useful for detecting structural defects before functional decline. This provides insight into early events of ATF6-related retinal vulnerability and a framework for future studies on rootlet-associated proteins and ciliary organization.

## Acknowledgments

Research results reported in this chapter were supported by NIH awards R01NS088485 and P30EY026877, VA Merits I01RX0002340, and P20 GM144230.

## Notes

### Competing Interest Statement

The authors have declared no competing interest.

